# Abscisic acid binds to an *Arabidopsis thaliana* phosphodiesterase and tunes its activity

**DOI:** 10.64898/2026.02.11.705244

**Authors:** Mateusz Kwiatkowski, Anna Kozakiewicz-Piekarz, Chuyun Bi, Aloysius Wong, Krzysztof Jaworski, Helen Irving, Chris Gehring

## Abstract

Growing evidence suggests that plant proteomes contain numerous proteins that specifically bind abscisic acid (ABA). Many of them are complex multidomain proteins where specific ABA-binding can cause biochemical and physiological changes. Here we show that the *Arabidopsis thaliana* K^+^ transporter AtKUP5 contains both a functional cytoplasmic N-terminal adenylate cyclase (AC) enabling the synthesis of 3’,5’-cAMP from ATP and a C-terminal phosphodiesterase (PDE) that hydrolyses 3’,5’-cAMP to 5’-AMP. We found that ABA binds in a ligand-specific manner to the catalytic center of the PDE thereby causing a reduction of 3’,5’-cAMP hydrolysis *in vitro*. The hydrolytic activity of the PDE is ABA concentration-dependent, biphasic and requires the presence of an intact ABA-binding site similar to the one in the canonical Pyrabactin resistance 1/PYR-like/Abscisic acid receptors, with V_max_ of 1.19 pmole min□^1^ μg□^1^ in the absence of ABA, increasing to 1.58 pmole min□^1^ μg□^1^ at 2 nM ABA, and decreasing to 0.75 pmole min□^1^ μg□^1^ at 50 nM ABA. These findings are therefore consistent with a direct role of ABA in PDE activity modulations and form a functional link between 3’,5’-cAMP signaling and K^+^ flux. Furthermore, we predict that a growing number of such receptor-like proteins that specifically and directly interact with ABA will be discovered thereby uncovering complex and ancient layers of signaling and metabolic regulation.

## Introduction

Abscisic acid (ABA) is a key regulator of many plant responses including seed maturation, germination, stomatal guard cell movement, dormancy, growth and stress responses (Ng et al. 2014). When an ABA receptor was first identified in 2009, the downstream signaling pathway was recognised as the key to ABA-dependent responses (Nishimura et al. 2010; Park et al. 2009). The model predicts that ABA operates as ligand for PYR/PYL/RCAR receptors (Pyrabactin resistance 1/PYR-like/Abscisic acid receptor). PYR/PYL/RCAR receptors are members of the family of ATPase transport proteins. Binding of ABA to the receptor inhibits protein phosphatases 2C (PP2C) and triggers ABA-dependent downstream signaling including SNF-related serine/threonine-protein kinase (SnRK2) activation. SnRK2 kinases are the primary kinases implicated in the control of stress responses. Transcription factors and other effector proteins are phosphorylated when SnRK2 kinases are activated. ABF (ABRE-binding factor) and MYB, for instance MYB2, 41, 44 and 96, are transcription factors that are activated by ABA, and they then enable ABA-dependent transcriptional programs (Fujita et al. 2011; Munemasa et al. 2015; Ng et al. 2014; Weiner et al. 2010).

An important question that arises is whether or not the PYR/PYL/RCAR receptor can fully account for all ABA-dependent processes. So far, four examples suggest that they may not. The first notable case is the relationship between ABA and mitochondrial adenine nucleotide translocators (ANT) (Kharenko et al. 2011). Through ATP/ADP exchange across the inner membrane of the mitochondria, ANTs help control the rate of ATP synthesis, an essential cellular process since cellular metabolism is inhibited when ANT activity declines (Neuhaus et al. 1997). Notably ABA can bind to ANT transporters thereby modifying their activity (Kharenko et al. 2011). Both native isolated spinach mitochondria and recombinant *A. thaliana* ANT2-containing proteoliposomes showed ANT-dependent ABA uptake with increasing administered ABA concentrations found to stimulate increased ATPase activity in mitochondrial extracts from spinach (Kharenko et al. 2011). A second example is the ABA interaction with Rubisco (Ribulose-1,5-Bisphosphate Carboxylase/Oxygenase). By determining the dissociation constant at 47 nM, it was possible to validate the high affinity of ABA for Rubisco and to identify possible binding sites for ABA using chemical proteomics methods and structural investigations (Galka et al. 2015). Furthermore, it was also suggested that ABA may function as a negative regulator of Rubisco activation since ABA causes a suppression of Rubisco catalytic activity but a considerably higher inhibition of enzyme activation. These findings are consistent with direct ABA-Rubisco interactions where, under specific conditions; ABA can modulate Rubisco activity presumably to ensure plant homeostasis (Galka et al. 2015). A third example is the direct and specific binding of ABA to the Guard Cell Outward-Rectifying Potassium Channel (GORK) which mediates K^+^ efflux from guard cells thereby causing stomatal closure. Whole-cell current–voltage experiments on HEK293 cells transfected with GORK showed that ABA directly increases K□efflux in a non-homologous system and hence in the absence of the PYR/PYL/RCAR receptor (Ooi et al. 2017). It is noteworthy that alterations of critical amino acids in the GORK ABA-binding site, results in an attenuation of the response to ABA-dependent response. A fourth and most recent example is the nitrate level sensing transceptor NRT1.1 which also interacts with ABA in a specific manner thereby integrating the plant nutrient status with ABA-dependent stress signalling (Ma et al. 2025; Wang and Tsay 2025).

Given that a conserved ABA-binding amino acid motif may identify candidate ABA-binding proteins other than GORK (Wong et al. 2022), we were interested to see if ABA-binding domains are present in complex regulatory proteins and if so, if we could establish biochemical and physiological functions of such binding. Here we report that the majority of identified plant phosphodiesterases (PDEs) which break down cyclic mononucleotides, notably cAMP and cGMP, to AMP or GMP respectively, harbour ABA-binding motifs in their sequences. This may be indicative for a novel mechanism by which ABA modulates cAMP homeostasis and signaling. We also demonstrate that ABA specifically directly binds to and modulates the hydrolytic activity of a PDE that is part of the *Arabidopsis thaliana* K^+^-Uptake Permease (AtKUP5) and this conceivably makes AtKUP5 an ABA receptor. Finally, if we accept that PDEs can be considered as an “off” signal in cyclic mononucleotide signaling, then modifying PDE activity with ABA may constitute a novel signaling mode of ABA.

## Results

### ABA-binding sites are present in plant phosphodiesterases

Previously built databases of *Arabidopsis thaliana* PDEs (Kwiatkowski et al. 2022; Kwiatkowski et al. 2021) were scanned for the presence of an amino acid ABA-binding motif (Ooi et al. 2017; Wong et al. 2022) . This motif consists of four key amino acid residues that form a binding site (Fig. 1A) in previously identified and confirmed ABA-binding candidates. Of the 53 plant PDEs, 30 harbour an ABA binding motif often close to or overlapping the catalytic center of the PDE domain (Supplementary Table 1). We selected the *Arabidopsis thaliana* K^+^ uptake permease 5 (AtKUP5; At4g33530) to examine ABA effects on the PDE which is located in the C-terminal cytoplasmic domain. In AtKUP5 the PDE catalytic center (Y669 – E706) overlaps with the ABA-binding site (E657 – K685) (Fig. 1A, B). Further analysis of the AtKUP5 structure generated by AlphaFold (Jumper et al. 2021) revealed two possible ABA binding sites as ascertained by the presence of surface grooves or pockets that could spatially accommodate the ligand. Molecular docking showed that both binding sites could dock with ABA, and the ligand assumes binding poses that have close molecular distances with key amino acids in the motif, Y678 and E657. ABA docked with good binding affinities of -4.8 kcal/mol and -4.7 kcal/mol at the respective Y678 and E657 pockets. Since the site of E657 appears to overlap with cAMP binding site of PDE, we performed docking of 3’,5’-cAMP at the PDE domain to assess how ABA might interfere with the activity of PDE. 3’,5’-cAMP could dock at the PDE catalytic center with a good binding affinity of -5.3 kcal/mol. Notably, the presence of ABA at the site of E657 does not seem to cause steric hindrance but rather a tighter fit of 3’,5’-cAMP at the overlapping PDE domain in the manner of orthosteric regulation, thus hinting at a possible enhancement of cAMP hydrolysis (Fig. 1C).

**Figure 1.**
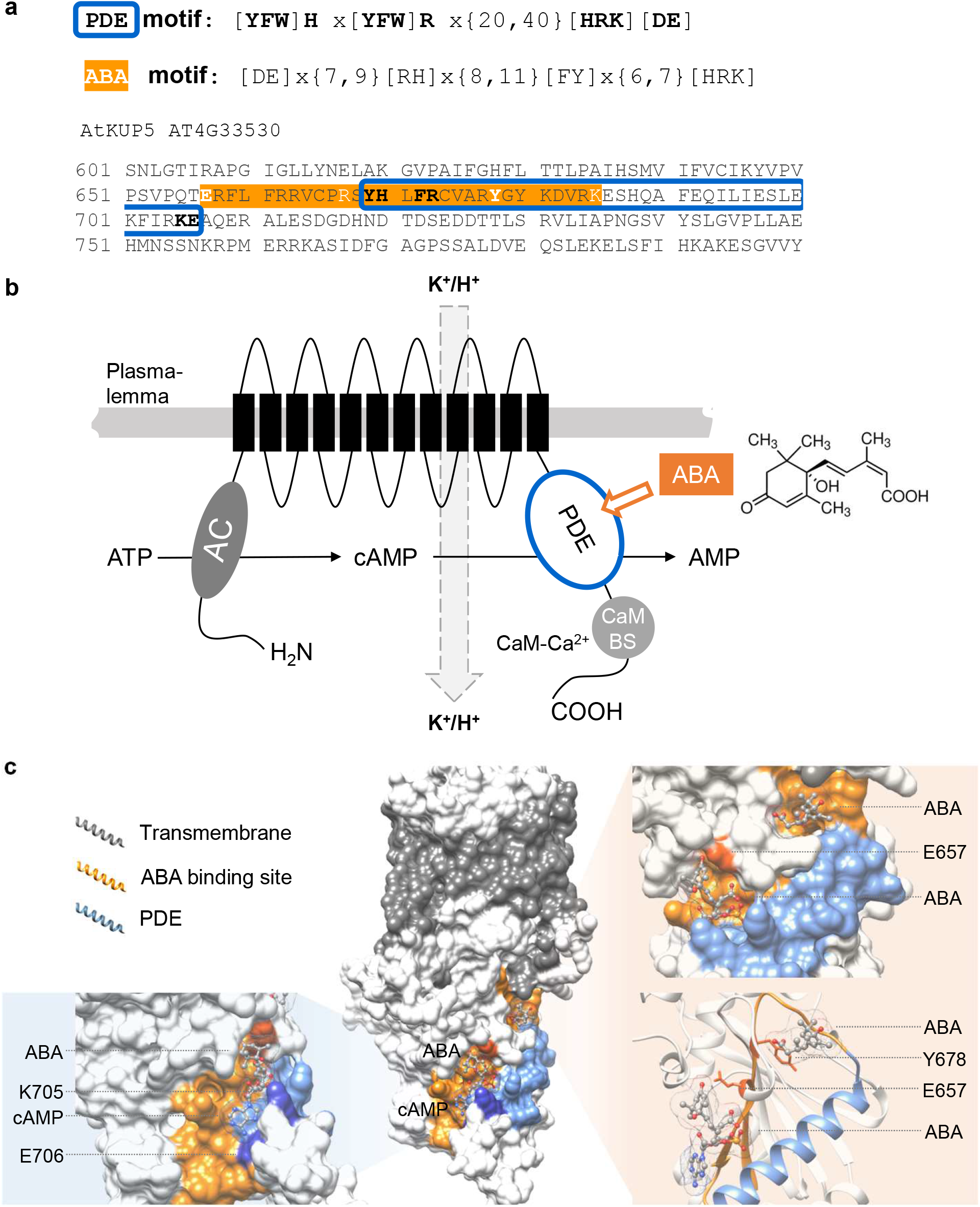
Sequence and structural analyses of AtKUP5 ABA binding site. **A** Amino acid sequence of AtKUP5, highlighting (orange) the ABA-binding site with the motif key amino acids bolded. Sequence of the PDE motif is framed in blue. **B** Architectural schematic of AtKUP5 for the dampening function of PDE in K^+^ transport regulation, illustrating the N-terminal AC center that is regulated by K^+^ ions to generate cAMP, a substrate for C-terminal PDE that is activated by the Ca^2+^-calmodulin complex or altered allosterically by ABA. **C** A three-dimensional model of AtKUP5 docked with ABA and cAMP. Molecular docking studies indicate that AtKUP5 could dock with ABA at two sites, forming interactions with key amino acids E657 and Y678 in the motif. The presence of ABA in the pocket of E657, may lead to a tighter fit of cAMP at the PDE domain. Structural assessment and image preparation were performed with AutoDock Vina (version 1.1.2) and UCSF Chimera. -

### ABA concentration-dependent effects on PDE activity

PDE catalytic activity was evaluated by enzymatic tests with 3’,5’-cAMP as the substrate in the presence or absence ABA. A sensitive liquid chromatography tandem mass spectrometry (LC-MS/MS) approach was used to identify and quantify the AMP products where we observe 5’-AMP formation, indicating hydrolysis of only the 3’5’-cAMP isoform (Fig. 2A). The results of the *in vitro* reaction showed that the hydrolytic activity of PDE changes depending on the concentration of ABA in the reaction mix (Fig. 2B). The PDE activity alters between concentrations of 2 to 50 nM ABA. The highest amount of cAMP hydrolysis occurs at 2 nM ABA, where the activity rises by > 30 % as compared to the control (absence of ABA) (Fig. 2B). Hydrolysis appears inhibited at ABA concentrations of 50 nM and above translating to a decrease in activity of about 30 % as compared to the control (Fig. 2B). The kinetic parameters of the reaction at the different concentrations of ABA indicates that ABA, even at 50 nM, raises the apparent substrate affinity for the enzyme as compared to the control reactions (K_*M*_ of 5.21 µM) (Fig. 2C). This is best demonstrated by the addition of 10 nM ABA, which reduces the K_*M*_ constant by more than twofold (K_*M*_ of 2.18 µM). The maximum reaction rate increases only in the presence of 2 nM ABA. Further increases in ABA concentration result in a > 35 % reduction in the apparent V_max_. While Ca^2^□-calmodulin (Ca^2+^/CaM) complexes enhance PDE activity of AtKUP5 as previously reported (Kwiatkowski et al. 2021), the addition of ABA results in an inhibition of PDE, independent of Ca^2^□/CaM *in vitro* (Supplementary Fig. 1).

**Figure 2.**
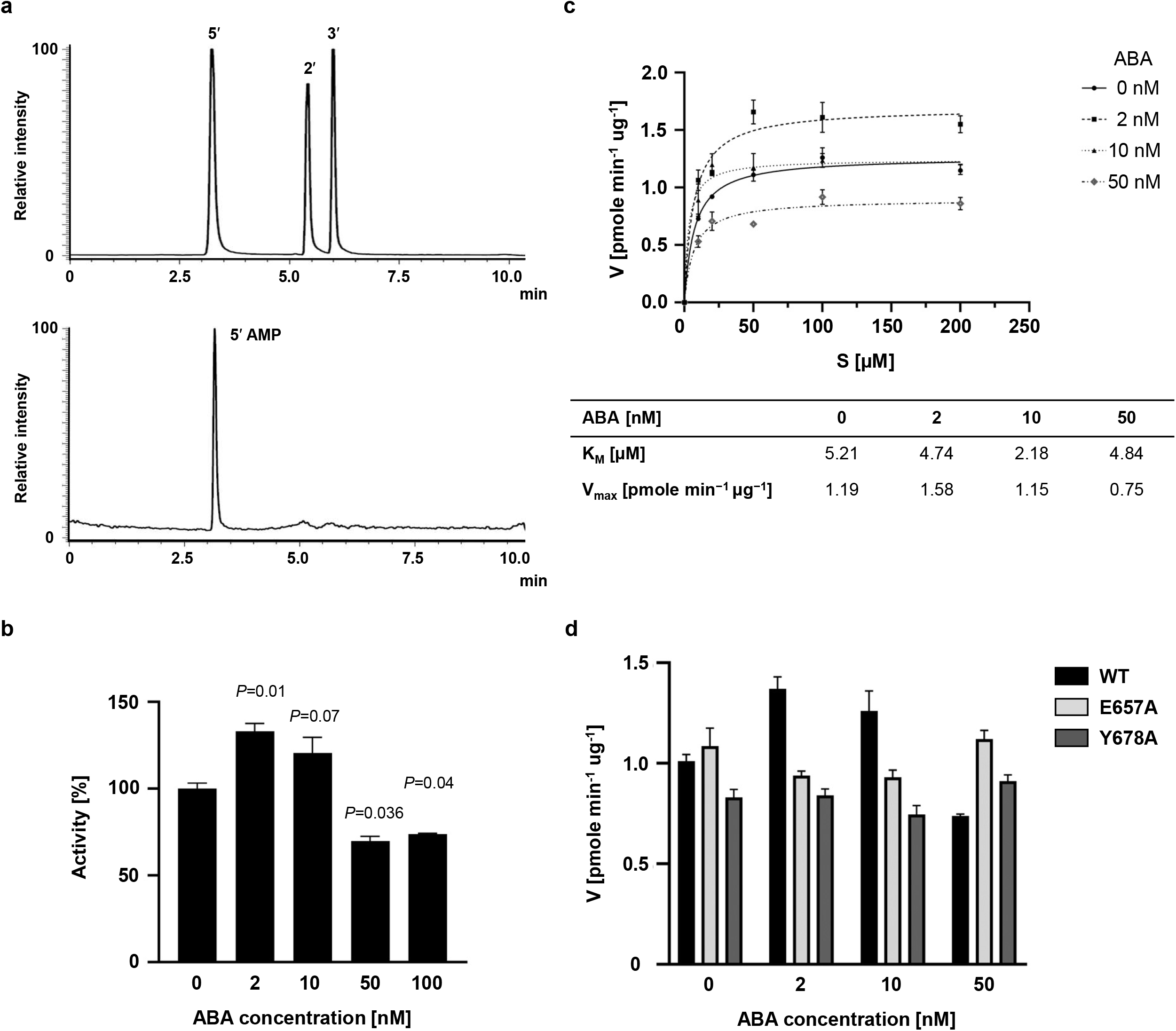
Effect of abscisic acid on the modulation of PDE activity. **A** Substrate specificity of AtKUP5 PDE. The upper panel shows the retention time of the separation of a standard of the three AMP isoforms. The lower panel shows the result of the separation of the PDE reaction using the substrates 2’3’-cAMP and 3’5’-cAMP. Only the 5’-AMP isoform is visible in the chromatogram, indicating hydrolysis of only the 3’5’-cAMP isoform. **B** PDE activity of AtKUP5 at different ABA concentrations. Data are mean values (n = 3) and error bars show the standard error of the mean. Statistical analysis was performed by Student’s t-test for independent samples with different variances. P < 0.05 indicates that the difference between groups is statistically significant. **C** Michaelis-Menten plot for PDE activity towards cAMP substrate in the presence of different ABA concentration. The inset shows table with a comparison of the kinetic parameters of the reaction with ABA at concentrations ranging from 0 to 50 nM. Data are mean values (n = 3) and error bars show the standard error of the mean. **D** Effect of single point mutations E657A and Y678A on PDE activity in the presence of different ABA concentrations. Data are mean values (n = 3) and error bars show the standard error of the mean.

Since increases in ABA inhibit PDE activity *in vitro*, the level of endogenous 3’5’-cAMP was determined in *E. coli* induced to express AtKUP5 and consequently more PDE is present in cells (Supplementary Fig. 2). AtKUP5 expression leads to a decrease in endogenous cAMP levels compared to the control, which is more enhanced in the presence of 2 nM ABA versus 100nM ABA.

### Mutation of amino acids in the ABA-binding domain affect interactions with PDE

The 3D model and docking analyses revealed that ABA may dock in two pockets with varying affinities, with the interacting amino acids being Y678 and E657. Therefore, we substituted both these amino acids with alanine obtaining AtKUP5^E657A^ and AtKUP5^Y678A^. The mutants were tested for PDE activity in the presence or absence of ABA (2 to 50 nM). *In vitro* reactions showed that the addition of ABA, unlike in the case of the wild type AtKUP5, did not significantly affect the course of the reaction, consistent with the predicted essential functions of E657 and Y678 in ABA binding (Fig. 2D). Expression of the AtKUP5^Y678A^ mutant also decreases endogenous cAMP, but, unlike in wild type AtKUP5, no fluctuations were seen in the level of cAMP hydrolysis after the addition of different concentrations of ABA (Supplementary Fig. 2).

To further characterise the protein-ABA interaction, fluorescence spectroscopy was employed. ABA (0 nM to 16 nM) was added to both the wildtype and the mutated proteins (Fig. 3). AtKUP5 exhibits a clear peak of fluorescence emission at 333 nm originating from tryptophan residues with an excitation of 280 nm. While the maximal emission wavelength remains constant, the intensity of the PDE fluorescence rises (or decreases in the case of the E657A mutant) as the ABA concentration increases (Fig 3A-C). However, in the mutants with altered ABA-binding pockets the increase is much smaller, as reflected by the calculated binding constants (Fig. 3D). The binding constant for AtKUP5 wildtype is 19.27 ± 3.94 [L•mol^-1^] x 10^8^, while for AtKUP5^E657A^ is 4.63 ± 1.44 [L•mol^-1^] x 10^8^ and for AtKUP5^Y678A^ is 2.03 ± 0.62 [L•mol^-1^] x 10^8^. This suggests that ABA-binding to the AtKUP5^E657A^ mutant is reduced four-fold while the reduction of ABA-binding in the AtKUP5^E657A^ is almost ten-fold. ABA-specific binding is further supported as only residual binding with indole acetic acid (IAA) or jasmonic acid (JA) was detected (Supplementary Fig. 3). ABA binds to AtKUP5 with a substantially higher affinity (10□ L/mol) than JA and IAA, for which association constants in the 10□ L/mol range were obtained.

**Figure 3.**
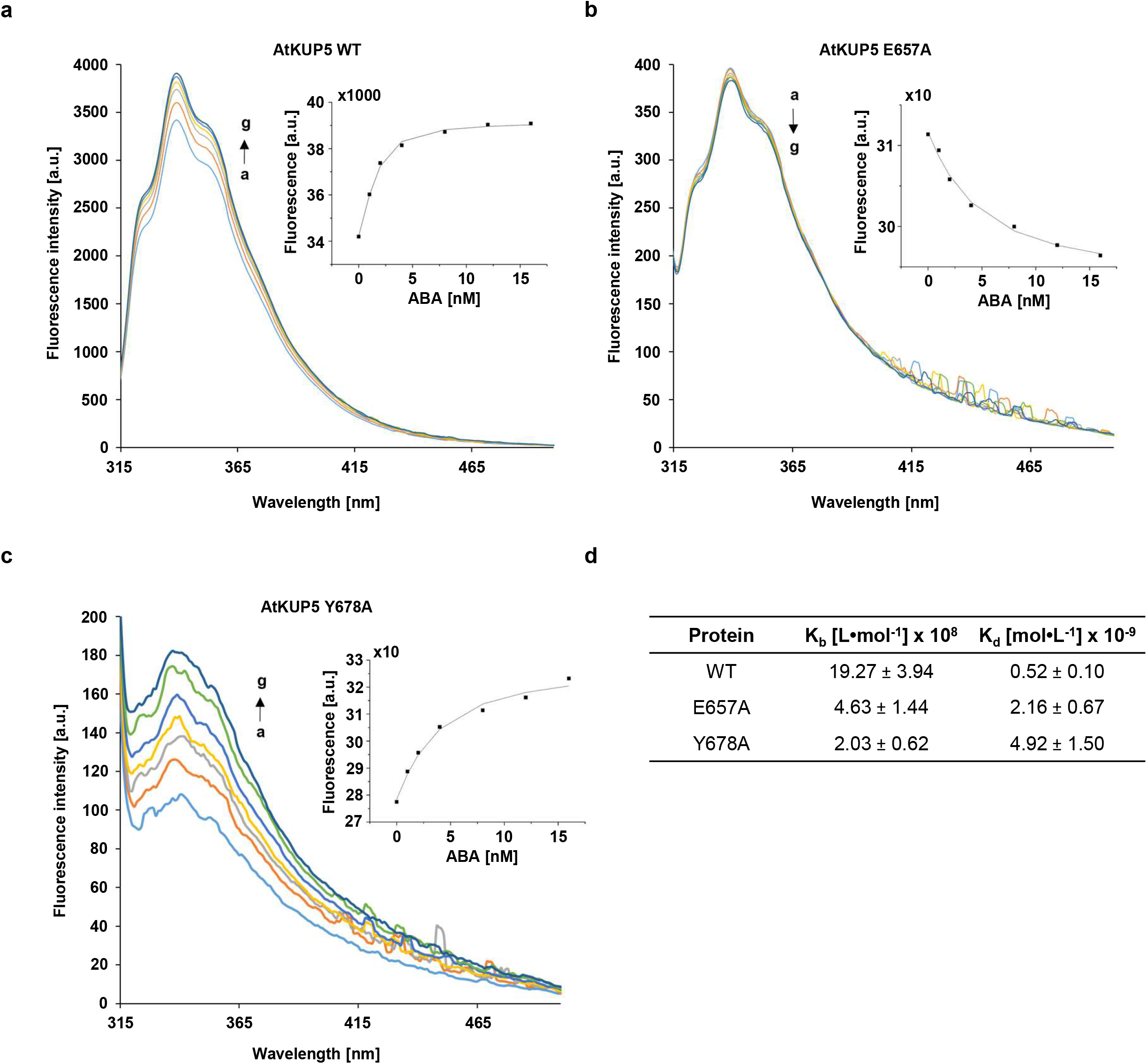
The fluorescence spectra of AtKUP5 wildtype and mutants in the presence of ABA. **A** The fluorescence enhancement spectra of AtKUP5. **B** The fluorescence enhancement spectra of AtKUP5^Y678A^. **C** The fluorescence quenching spectra of AtKUP5^E657A^. The concentration of recombinant protein was 2 nM and treatments were at 22°C. The concentration of ABA ranged from a to g were 0, 1, 2, 4, 8, 12, and 16 nM, respectively. **D** Comparison of kinetic parameters of interactions between ABA and AtKUP5 wildtype and mutants, K_b_ – binding constant, K_d_ – dissociation constant.

## Discussion

The discovery of a conserved ABA-binding amino acid signature in many proteins suggested that ABA-signalling might be more complex and intricate than hitherto suspected. The first indication of widespread ABA-binding to target proteins came when GORK was shown to bind ABA and that this binding is causative for modulations of the K^+^-flux through the channel (Ooi et al. 2017). The amino acid motif appears to be highly conserved and present not just in the canonical ABA receptor but also but also in > 200 candidate proteins in the *Arabidopsis thaliana* proteome (Wong et al. 2022). Among these are a number of PDEs (Supplementary Table 1) and proteins with dual or multiple activities (e.g. adenylate cyclase (AC) and PDE) (Kwiatkowski et al. 2022; Turek et al. 2024). This may suggest that by affecting either the AC or PDE activity or both, such dual activity proteins can modify 3’,5’-cAMP homeostasis and signaling and hence operate as 3’,5’-cAMP tuners. Such dual or indeed multifunctional proteins are also referred to as moonlighting proteins (Irving et al. 2012; Wong et al. 2015) that are increasingly recognised as critical signaling modules that can integrate multiple second messengers such as Ca^2+^ and 3’,5’-cAMP and 3’,5’-cGMP (Irving et al. 2012; Muleya et al. 2014; Świeżawska-Boniecka et al. 2021).

Here we show that AtKUP5 which contains functional AC (Al-Younis et al. 2018a) and PDE catalytic centers (Kwiatkowski et al. 2021) also harbours an ABA-binding motif close to the PDE center. Structural analysis combined with molecular docking studies revealed that the ABA-binding motif spans multiple pockets within the PDE catalytic center and points to essential roles for E657 and Y678 in ABA-PDE interaction. Since substitution of these residues with alanine resulted in a significant loss of ABA-dependent modulation of PDE activity (Fig. 2) as well as interactions (Fig. 3), the ABA pockets might therefore confer allosteric regulation or in the case of the pocket at E657 orthosteric regulation, where the presence of ABA causes a tighter fit of cAMP that favours hydrolysis (Fig. 1C). High ABA concentrations could however saturate the cAMP binding site blocking its interaction with the PDE, and this might account for the biphasic kinetic behaviour observed in our study. Taken together, these findings not only validate AtKUP5 as an ABA-responsive protein but also demonstrate that the ABA-binding motif is an effective tool for the discovery of ABA-binding candidates.

*In vitro* experiments demonstrated that 3’,5’-cAMP-specific PDE activity is modulated by ABA in a biphasic manner. At low ABA concentrations (2–10 nM), there is a notable increase in hydrolytic activity, whereas higher concentrations (50 nM and above) there is inhibition (Fig. 2A). It is conceivable that ABA can prevent signal attenuation at higher concentrations to maintain cellular sensitivity to stimuli possibly by preventing cAMP binding to the active site. Alternatively, ABA may promote cAMP hydrolysis at low concentrations to tune downstream reactions. This pattern is consistent with known hormone dose–response relationships, where low concentrations promote signaling or activity, while higher concentrations trigger negative feedback or receptor saturation effects (Umezawa et al. 2010). At low nanomolar concentrations (2–10 nM), ABA likely promotes the active conformation of the PDE complex. Such concentrations are within the physiological range of ABA reported in well-watered plants (Wilkinson and Davies 1997), and may serve as a signal to modulate enzyme activity under basal conditions. The decline in V_max_ observed at higher ABA concentrations reflects a shift toward an inhibitory mechanism which may result from allosteric inhibition or conformational destabilization of the catalytic domain. High ABA levels are known to trigger feedback loops and downstream signaling cascades that modulate the sensitivity and output of ABA-responsive proteins (Fujita et al. 2011).

Biphasic responses are not entirely uncommon (Agathokleous et al. 2019; Caraty et al. 1989; Li et al. 2022; Li et al. 2017). Characteristics of dose responses have recently been reviewed and revaluated (Agathokleous et al. 2019). It appears that perhaps contrary to expectations, linear dose–responses are not the default response to hormones but the biphasic dose–response is. This phenomenon is termed hormesis and may point the beneficial effects of basal level stress for optimum health. Hormesis has been shown to occur in many species and it appears that the low stress responses are a precondition or priming point which in turn is essential to survive otherwise lethal massive stresses (Ling et al. 2018).

There is now a growing body of evidence for ABA-protein interactions that function outside of the classic PYR/PYL/RCAR-PP2C-SnRK2 signaling pathway and the direct ABA effect on AtKUP5 PDE function shown here is a point in case. Direct effects of ABA on protein function was also previously reported in other proteins, namely Rubisco (Galka et al. 2015) and GORK (Ooi et al. 2017). In AtKUP5 and GORK the ABA effect could be attributed to the presence of conserved and specific ABA-binding amino acid motif. In contrast, the effect of ABA on Rubisco in *Pisum sativum* did not appear to be dependent on the presence of the strict [DE].{7,9}[RH].{8,11}[FY].{6,7}[HRK] motif. Crystallographic analysis suggested that ABA might interact with the large subunit of Rubisco and at a site distinct from the active site, a classic example of allosteric modulation. Interestingly, it was also suggested that a possible interaction could occur with arginine, glutamic acid, and tyrosine, hence the residues that are part of the ABA-binding motif described here. Although it has been shown that ABA binds to the non-activated form of the enzyme, ABA might still act as an inhibitor of Rubisco activation. However, only mutagenesis of the key amino acids indicated in the two sites might eventually elucidate the physical nature of these ABA effects. Incidentally, the ABA-binding motif [DE].{7,9}[RH].{8,11}[FY].{6,7}[HRK] is present in the Rubisco activase (At2g39730, Supplementary Fig. 4). The Rubisco activase removes the inhibition of phosphorylated compounds that bind to both carbamylated and non-carbamylated Rubisco active sites thereby modulating Rubisco activity in response to temperature and light variations. These inhibitory substances, such as RuBP, are removed by Rubisco activase using energy from ATP hydrolysis, restoring Rubisco catalytic activity (Andersson 2008). The ABA-binding motif in this enzyme is located in the region spanning the Box VII′ and Sensor 2 domains at position 343–368, i.e. in the region responsible for substrate recognition and conformational remodeling of Rubisco (Portis et al. 2008). Given that ABA has been demonstrated to interact with Rubisco, it is conceivable that Rubisco activase and ABA cooperate and that ABA functions as an effector molecule that can either stabilize or destabilize the interaction between Rubisco and Rubisco activase. Additionally, ABA and Rubisco activase may interact to decrease Rubisco activity when photosynthetic efficiency is impaired, directing resources and energy toward stress responses.

The role of cyclic nucleotide monophosphates (cNMPs) in ABA signalling is well established (Al-Younis et al. 2021; Liang et al. 2022), however, most attention is paid to the processes in which ABA participates in signalling rather than to the description of specific signalling components and their mechanisms of action (Świeżawska-Boniecka and Szmidt-Jaworska 2023). There are known ABA-downstream responses that require cAMP or cGMP e.g. stomatal movement, seed germination or the production of isoflavones under UV-B stress in soybean (Jiao and Gu 2019; Jin and Wu 1999; Liu et al. 2023). It is noteworthy that ABA and the recently identified nucleotide cyclases are in some instances closely linked. One such case is the essential chloroplast enzyme involved in ABA production, the *A. thaliana* 9-cis-epoxycarotenoid dioxygenase 3 (AtNCED3). AtNCED3 is both essential for the synthesis of ABA in the carotenoid pathway and synthesis of cAMP since it functions as an AC (Al-Younis et al. 2021). The maize ZmRPP13-like protein 3 (ZmRPP13-LK3) is a heat-induced potential disease-resistance protein found in mitochondria that also functions as an AC (Yang et al. 2021). Under heat stress, ABA increases both cAMP levels and the transcriptional activity of the ZmRPP13-LK3. *In vivo* investigations revealed that cAMP may bind to the ZmABC2 that operates as a potential cAMP exporter. It appears that cAMP contributes to the resilience of maize to heat stress and functions as a downstream component of the ABA-mediated stress response. Direct interactions between ABA and the K^+^ transporter AtKUP5 are consistent with mechanisms that link ABA and cNMPs and ion homeostasis. The control of cellular ion balance is influenced by ABA (MacRobbie 2000) and AtKUP5 might well function as cAMP-dependent K^+^-flux sensor (Al-Younis et al. 2018) where ABA modulates cAMP generation which in turn is directly dependent on K^+^ flux. Moreover, members of the KT/HAK/KUP potassium transporter family show stress□regulated expression and are implicated in abiotic stress responses, including via modulation by ABA (Yang et al. 2020). ABA, as a central stress hormone, triggers cytosolic Ca^2^□elevations, which, together with calmodulin, is an activator of AtKUP5 (Kwiatkowski et al. 2021). While Ca^2^□/CaM enhances PDE activity, the addition of ABA inhibits PDE, independent of Ca^2^□/CaM (Supplementary Fig. 1). This suggests that ABA functions as an upstream stress signal that suppresses cAMP degradation, thereby modulating cyclic mononucleotide signaling under stress conditions. In addition, AtKUP5 function *in planta* has been studied in knockout lines and overexpression lines where it was shown that deletion of AtKUP5 causes both decreased K^+^ uptake and reduced levels of cellular 3’,5’-cAMP as well significantly impaired root elongation (Al-Younis, 2018b). To elucidate the specific effects of ABA on AtKUP5-dependent K^+^ uptake and its dependence on the AtKUP5-PDE, mutants with altered ABA-binding sites will have to be tested. Such mutant analyses will also allow testing of the hypothesis that (stress-induced) ABA surges inhibit PDE activity and hence cause increases of cAMP, and this in the wildtype only. This hypothesis seems to be valid for *in vivo* tests using *E. coli*, where in the presence of various ABA concentrations we determined the level of endogenous cAMP in lines overexpressing AtKUP5 and the ABA-binding mutant AtKUP5^Y678A^ (Supplementary Fig. 2). In the case of AtKUP5 protein overexpression, and consequently an increase in the amount of PDE in cells, a decrease in the endogenous cAMP level is observed compared to the control, which is even more enhanced when ABA is added at a concentration of 2 nM. Simulation of stress by incubating cells with 100 nM ABA inhibits PDE activity, equalizing the cAMP level to the initial conditions. In turn, overexpression of the ABA-binding PDE mutant results in a decrease in the level of endogenous cAMP during induction, but we did not observe fluctuations in the level of cAMP hydrolysis after the addition of different concentrations of ABA, as was the case with the AtKUP5 protein, thus reflecting the performed *in vitro* assays.

In summary, we present further support for the notion that, first, specific ABA-binding can occur in a number of proteins and that this binding can exert distinctive biochemical and physiological effects on the target proteins. Given that ABA does occur in lower plant, fungi and animals (Gharib et al. 2024; Ng et al. 2014), it makes this hormone an ancient regulator of numerous biological responses and processes.

## Materials and Methods

### Structural analysis of ABA binding site in AtKUP5

The structural prediction and docking were essentially conducted as detailed in (Zhou et al. 2021). The AtKUP5 model was obtained from AlphaFold (Jumper et al. 2021), available at https://alphafold.ebi.ac.uk/entry/Q8LPL8. The ABA and PDE binding sites were evaluated based on the presence of surface grooves or pockets that could spatially accommodate the ligand. Docking studies were conducted with KUP5 as the “receptor” and ABA or cAMP as the “ligand”. The generated binding affinities, ligand binding poses, molecular distances and interactions with key amino acids at the ABA binding sites were assessed. Molecular docking studies were conducted using AutoDock Vina (version 1.1.2) (Trott and Olson 2010) and structural visualization and image preparation were performed using UCSF Chimera (Pettersen et al. 2004).

### PDE biochemical assay and LC-MS/MS analysis

Recombinant PDE domain of AtKUP5 was prepared as described in the Supplementary methods. PDE *in vitro* activity was determined by using LC-MS/MS to determine the rate of AMP formation. The reaction mixture contained: 3 mM Tris-HCl (pH 8.0), 0.01 to 0.2 mM cAMP, 0.1% (v/v) 2-mercaptoethanol, 5 µg of recombinant protein (supplementary methods), 0.5 mM MgCl_2_ and MnCl_2_, and 2 to 100 nM ABA. Samples were incubated at 37 °C for 20 min. The enzyme reaction was terminated by incubation at 100 °C for 5 min and the samples were centrifuged at 13,200 × g for 10 min.

LC-MS/MS experiments were performed using the Nexera UHPLC and LCMS-8045 integrated systems (Shimadzu Corporation, Kyoto, Japan). The ionization source parameters were optimized in positive ESI mode using pure AMP dissolved in HPLC grade water (Sigma, St. Louis, MO, USA). The samples were separated using a Discovery HS C18 column (100 × 2.1 mm, 5 µm, Sigma, St. Louis, MO, USA). A gradient of solvent A (0.05 % (v/v) formic acid with 5-mM ammonium formate) and solvent B (100 % (v/v) acetonitrile) was applied over 3 min: B: 0–5%, followed by washing and conditioning of the column with a flow rate of 0.4 mL/min. The interface voltage was set at 4.0 kV for positive (ES+) electrospray. Data acquisition and analysis were made with the LabSolutions workstation for LCMS-8045.

### Fluorescence and interaction studies

The fluorescence spectra of proteins in the absence and presence of ABA were performed on a JASCO FP-8300 spectrofluorometer with a 10 mm quartz cell (Hellma Analytics, Müllheim, Germany). Measurements were recorded in the range of 300–600 nm after excitation at λ = 280 nm at 22 °C. The stock solutions of ABA, IAA and JA were prepared in phosphate buffered saline (PBS). The samples were prepared in 1.5 mL Eppendorf tubes, and they contained proteins (wild type, E657A, and Y678A) at a concentration of 2 nM without or with ABA at the following concentrations: 1, 2, 4, 8, 12, and 16 nM, and PBS. The fluorescence data were fitted by applying a nonlinear least-squares regression using OriginPro software Version 2016 (OriginLab Corporation, Northampton, MA, USA).

## Author contributions

C.G. conceived of the project, C.B. and A.W. did the structure modelling, M.K. and A.K.-P. performed the experiments and all authors contributed to the data analyses and writing of the manuscript.

## Supplementary Data

Supplementary data is available in the online file.

## Data Availability

The data underlying this article are available in the article and in its online supplementary material.

