## Supplementary file for "Abscisic acid binds to an *Arabidopsis thaliana* phosphodiesterase and tunes its activity"

The Following pages include:

- Supplementary Table 1. List of *A. thaliana* PDE-ABA-binding candidates
- Supplementary Fig. 1. Effect of abscisic acid on AtKUP5 activity in the presence of an active calmodulin complex.
- Supplementary Fig 2. *In vivo* analysis of ABA-dependent changes in intracellular cAMP levels in *E. coli* overexpressing AtKUP5.
- Supplementary Fig. 3. The fluorescence quenching spectra of AtKUP5 in the presence of (A) indole-3-acetic acid (IAA) and (B) jasmonic acid (JA).
- Supplementary Fig. 4. Alignment of the Rubisco activase derived from *A. thaliana* (At2g39730) and *P. sativum* (XP_050904391.1)
- Supplementary Methods

**Supplementary Table 1. List of *A. thaliana* PDE-ABA-binding candidates**

Search patterns:

PDE1: [YFW] H x [YFW] R x {20,40} [HRK] [DE]

PDE2: [AYFW] H x [LEYFW] R x {20,40} [HRK] [DE] x {60,90} R x {3} [YFW]

ABA: [DE] x{7,9}[RH] x{8,11}[FY] x{6,7}[HRK]

Dataset searched:	*Arabidopsis thaliana* proteome ID UP000006548

| Gene ID | ABA site | PDE site | Description |
| --- | --- | --- | --- |
| AT1G02120 | 381-410 | 430-544 | VASCULAR ASSOCIATED DEATH1 |
| AT1G03740 | 121-151 | 86-213 | Protein kinase superfamily protein |
| AT1G06950 | 705-735 | 593-721 | ARABIDOPSIS THALIANA TRANSLOCON AT THE INNER ENVELOPE MEMBRANE OF CHLOROPLASTS 110 |
| AT2G32000 | 42-69 | 95-210 | DNA topoisomerase, type IA |
| AT2G35110 | 351-377 | 308-441 | GNARLED, NCK-ASSOCIATED PROTEIN 1 |
| AT4G02430 | 143-169 | 78-191 | Serine/Arginine-Rich Protein Splicing Factor 34b |
| AT4G28080 | 53-77 | 109-203 | REDUCED CHLOROPLAST COVERAGE 2 |
| AT4G32180 | 103-131 | 588-705 | Pantothenate kinase 2 |
| AT5G10290 | 335-360 | 373-528 | Leucine-rich repeat transmembrane protein kinase family |
| AT5G51540 | 29-55 | 148-263 | Mitochondrial ATP-independent protease |
| AT1G77270 | 529-555 | 638-674 | Hypothetical protein |
| AT3G57200 | 149-179 | 334-367 | Glycosyltransferase-like protein |
| AT4G33530 | 657-686 | 669-706 | KUP5 potassium transporter |
| AT5G09400 | 656-685 | 668-705 | KUP7 potassium transporter |
| AT5G21160 | 758-785 | 781-815 | AtLARP1a, La related protein 1a |
| AT5G36930 | 168-194 | 355-399 | Disease resistance protein (TIR-NBS-LRR class) |
| AT1G17330 | 162-188 | 27-64 | Metal-dependent phosphohydrolase |
| AT3G17550 | 17-43 | 226-263 | Haloacid dehalogenase-like hydrolase (HAD) superfamily protein; |
| AT4G20760 | 234-262 | 208-234 | NAD(P)-binding Rossmann-fold superfamily protein |
| AT5G12080 | 98-124 | 436-462 | ATMSL10, MECHANOSENS. CHANNEL OF SMALL CONDUCTANCE |
| AT3G50140 | 351-378 | 152-178 | DUF247 |
| AT3G50190 | 302-329 | 135-161 | Transmembrane DUF247 |
| AT1G78240 | 267-294 | 116-145 | TSD2, TUMOROUS SHOOT DEVELOPMENT 2 |
| AT2G24740 | 34-65 | 634-670 | SDG21, SET DOMAIN GROUP 21 |
| AT1G71460 | 197-222 | 291-408 | Pentatricopeptide repeat (PPR-like) |
| AT2G03270 | 449-474 | 513-624 | DNA-binding protein |
| AT2G19380 | 99-127 | 302-417 | RNA recognition motif (RRM) containing protein |
| AT3G50280 | 50-79 | 266-383 | HXXXD-type acyl-transferase family protein |
| AT3G61800 | 54-82 | 359-496 | UVSSA ENTH/VHS protein |
| AT5G61980 | 467-492 | 223-320 | AGD1, ARF-GAP DOMAIN 1 |

**Supplementary Figure 1**

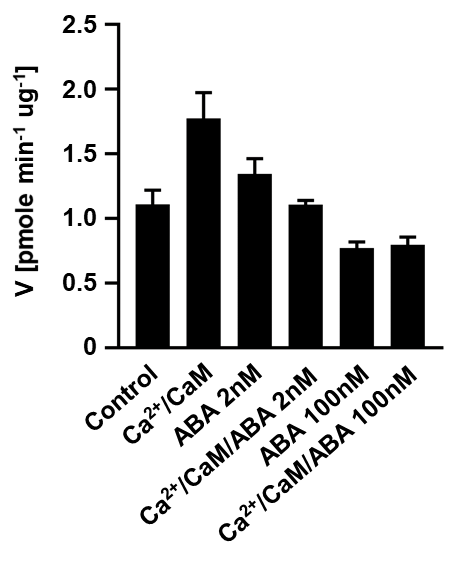

**Effect of abscisic acid on AtKUP5 activity in the presence of an active calmodulin complex.**

**Supplementary Figure 2**

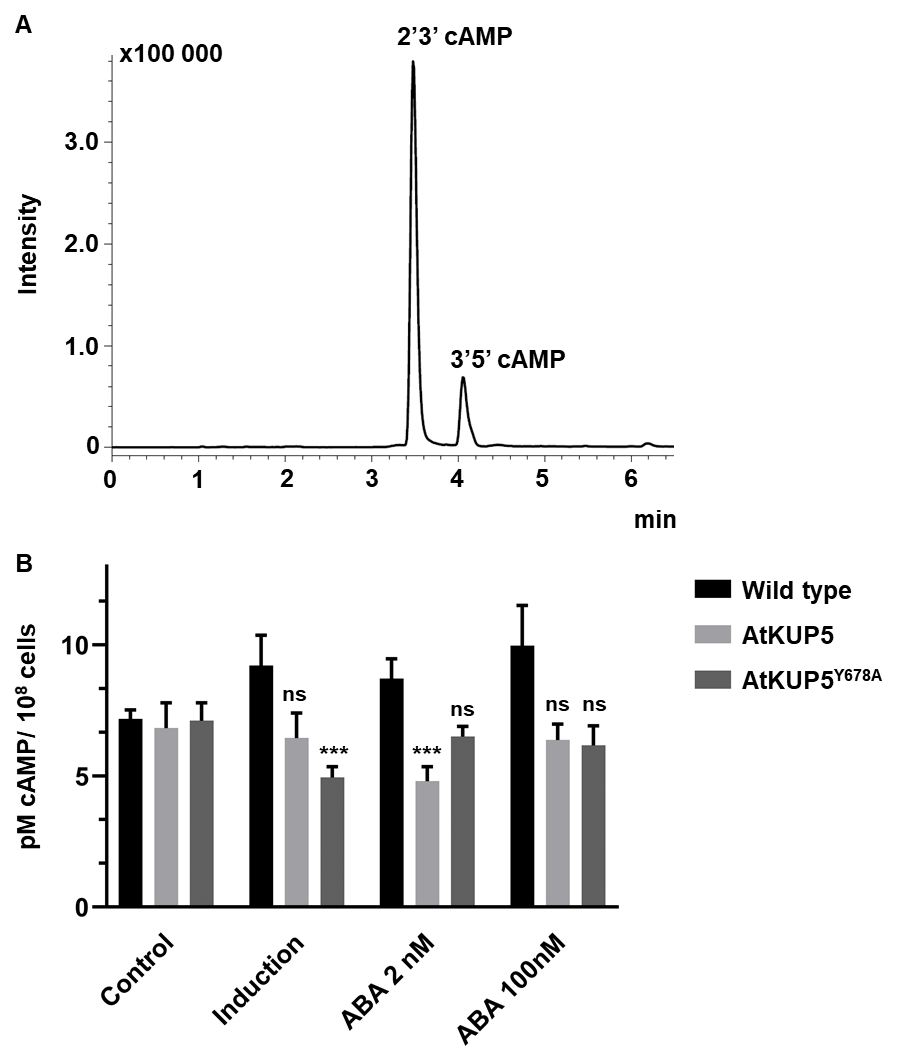

***In vivo* analysis of ABA-dependent changes in intracellular cAMP levels in *E. coli* overexpressing AtKUP5. (A)** Representative LC-MS chromatogram showing separation of intracellular cAMP isoforms, with 3’,5’-cAMP indicated. **(B)** Quantification of intracellular cAMP levels in *E. coli* cultures under control conditions, overexpression, and following ABA treatment in the wild type, AtKUP5 and AtKUP5^Y678A^ lines. Data are mean ± SEM (n=3). Significance was assessed by two‑way ANOVA with Tukey’s post‑hoc test. Symbols indicate significant difference compared to Wild-type Control: ns – not significant, *p < 0.05, **p < 0.01, ***p < 0.001.

**Supplementary Figure 3**

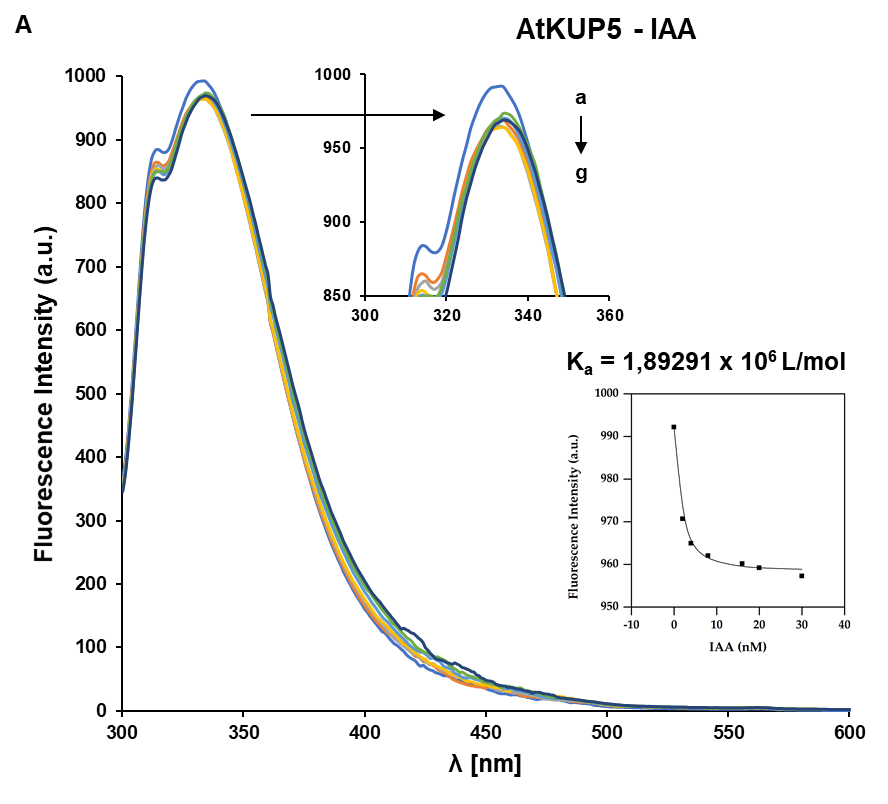

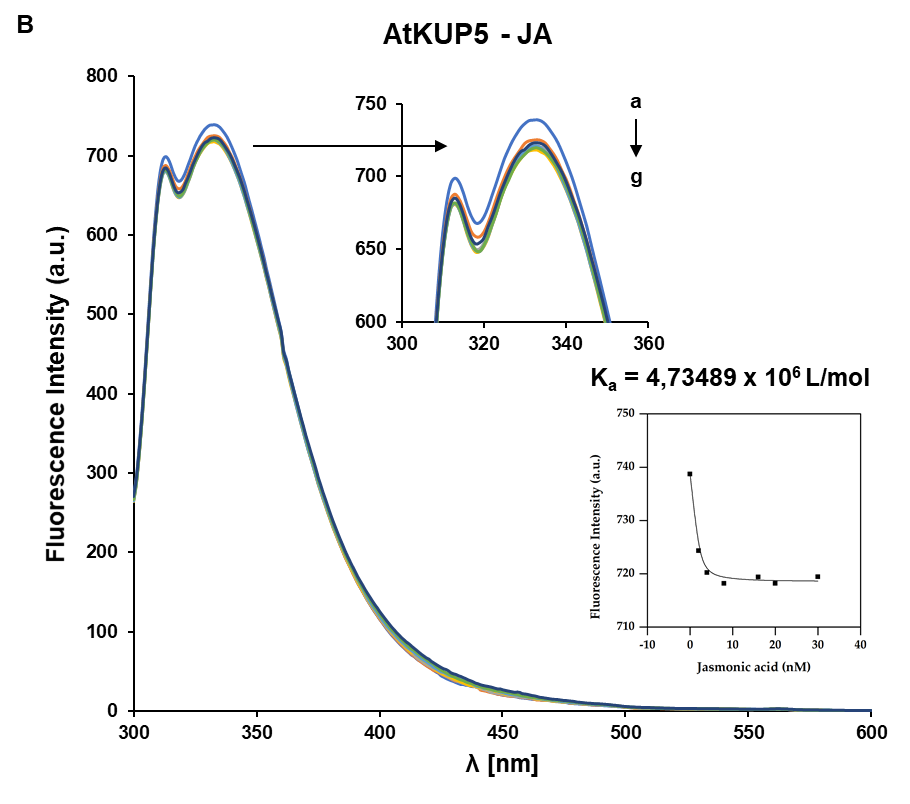

The fluorescence quenching spectra of AtKUP5 in the presence of **(A)** indole-3-acetic acid (IAA) and **(B)** jasmonic acid (JA). The concentration of recombinant protein was 2 nM. The concentrations of IAA or JA were from "a" to "g" were: 2, 4, 8, 16, 20, and 30 nM, respectively.

**Supplementary Figure 4**

```
P. sativum      GRMEKFYWAPTREDRIGVCKGIFRHDNVPEEEVVKIVDTFPGQSIDFFGALRARVYDDEV	359
A. thaliana     GRMEKFYWAPTREDRIGVCKGIFRTDKIKDEDIVTLVDQFPGQSIDFFGALRARVYDDEV	357
                ************************ *:: :*::*.:** *********************

P. sativum      RKWISGVGIEGIGKKLVNSKEGPPTFDQPKMTLEKLLEYGNMLVQEQENVKRVQLADKYL	419
A. thaliana     RKFVESLGVEKIGKRLVNSREGPPVFEQPEMTYEKLMEYGNMLVMEQENVKRVQLAETYL	417
                **::..:*:* ***:****:****.*:**:** ***:******* ***********:.**
```

Alignment of the Rubisco activase derived from *A. thaliana* (At2g39730) and *P. sativum* (XP_050904391.1) with ABA-binding motif highlighted in yellow.

**Supplementary methods**

*Expression vector construct and site directed mutagenesis of AtKUP5*

Total RNA was isolated from leaves and stalk of *A. thaliana* using RNeasy Plant Mini Kit (Qiagen, Hilden, Germany). The first-strand cDNA for RT-PCR was synthesized using GoScript™ Reverse Transcription System (Promega, Madison, WI, USA) following the manufacturer’s instructions. In order to construct the expression plasmids for the GST-AtKUP5^573-855^, a cDNA fragment encoding the PDE domain together with ABA binding site was amplified by RT-PCR using the specific primers:

- *AtKUP5^573-855^* (forward)

5ʹ-GGATCCCCAGGAATTCCCATGTTTGTTTGGAACTACGGGAG-3ʹ,

- *AtKUP5^573-855^* (reverse)

5ʹ-GATGCGGCCGCTCGAGAATCATACCATATAAGTCATTCCAACT-3ʹ,

The PCR reactions were performed with a cDNA as a template, forward and reverse primers and PrimeSTAR Max Polymerase (Takara Bio USA, Mountain View, CA, USA). The amplified DNA fragments were purified using Agarose-Out DNA Purification Kit (Eurx, Gdańsk, Poland). The amplified PCR products were cloned into pGEX-6P-2 vector (Cytiva, Uppsala, Sweden) in the *Eco*RI – *Xho*I restriction sites using an In-fusion Cloning kit (Takara Bio USA, Mountain View, CA, USA).

ABA binding site GST-AtKUP5^E657A^ and GST-AtKUP5^Y678A^ mutants were constructed by site directed mutagenesis using QuikChange II XL Site-Directed Mutagenesis Kit (Agilent, Cedar Creek, TX, USA). Specific primers used in the reaction were:

- *AtKUP5^E657A^* (forward)

5ʹ-GTCTGAAAAGAAATCTCGCGGTCTGAGGCACACT-3ʹ

- *AtKUP5^E657A^* (reverse)

5ʹ-AGTGTGCCTCAGACCGCGAGATTTCTTTTCAGAC-3ʹ

- *AtKUP5^Y678A^* (forward)

5ʹ-CGCACATCTTTGTATCCAGCCCTGGCTACACAGCGGAA-3ʹ

- *AtKUP5^Y678A^* (reverse)

5ʹ-TTCCGCTGTGTAGCCAGGGCTGGATACAAAGATGTGCG-3ʹ

*Expression and purification of the recombinant protein*

The resulting plasmids were introduced into the *E. coli* BL21(DE3) pLysS competent cells (Promega, Madison, WI, USA) in order to produce the fusion proteins with glutathione-S-transferase (GST) affinity tag. The transformants were grown in LB medium (250 mL) containing ampicillin (100 μg/mL) and 2 % glucose at 37 °C. Fusion protein expression was induced by adding isopropyl-ß-D-thiogalactopyranoside (IPTG) to a final concentration of 0.2 mM at OD_600_ = 0.6 and incubating the culture at 18 °C overnight. The bacteria were harvested by centrifugation and the pellet was suspended in lysis buffer (50 mM Tris-HCl pH 8.0, 150 mM NaCl, 5 mM EDTA, 5 mM EGTA, 1 % (v/v) Triton X-100, 1 mM PMSF, 0.2 mg/mL lysozyme) and disrupted by sonication. The cell extract was centrifuged at 18,000 × g for 30 minutes and the supernatant was loaded onto a glutathione-Sepharose 4B beads (Cytiva, Uppsala, Sweden). Afterwards the column was washed multiple times with buffer containing 50 mM Tris-HCl (pH 8.0) and 150 mM NaCl until the A_280_ was below 0.002 and the GST fusion protein was eluted with 10 mM glutathione in 50 mM Tris-HCl (pH 9.0). The homogeneity and purity of eluted protein fraction was analysed by SDS–PAGE electrophoresis (10 % gel) with the Coomassie Blue gel staining.

*cAMP isolation from bacteria cultures*

*E. coli* BL21(DE3) pLysS strains (Promega, Madison, WI, USA) containing the plasmid pGEX6p2 (WT), pGEX6p2-AtKUP5 and pGEX6p2-AtKUP5^Y678A^ were used to measure the amount of cAMP in response to abscisic acid. Aliquots (5 mL) of *E. coli* cultures grown in LB medium, with or without ABA, were induced with 1 mM IPTG at an OD_600_ of 0.6 to overexpress the corresponding protein variants. Cultures were incubated for 30 min at 30 °C, then chilled and centrifuged (4 °C, 3000 × g). Frozen cell pellets were suspended in 500 μL of ice-cold acetonitrile/methanol/water (2/2/1, v/v/v), then the cells were lysed by sonication on ice and subsequently centrifuged at 4°C at 8000 × g for 10 min. The lysate was and dried under vacuum and resuspended in 100 μL of water with 0.1% formic acid. The extracts were centrifuged at 12000 × g for 20 min at 4°C and transferred to HPLC vial.

The amount of cAMP was measured in 3 µL of the sample using LC-MS/MS. The analyses were performed using a Nexera UHPLC coupled with an LCMS-8045 integrated system (Shimadzu Corporation, Kyoto, Japan). The experiments were conducted in positive ESI mode, utilizing pure cAMP standards (Sigma, St. Louis, MO, USA) dissolved in HPLC-grade water. Cyclic AMP parent ion was set at m/z 330.00 [M + H]^+^ and the corresponding fragmented daughter ion at m/z 136.30 [M + H]^+^. The fragmented product ion was used for quantitation. Sample separation was carried out on a Supelco Ascentis Express C18 column (Sigma, St. Louis, MO, USA) employing a gradient of solvent B (2%-10%) over 8.5 minutes, with solvent A consisting of water containing 0.1% formic acid (v/v) and solvent B being 80% acetonitrile with 0.05% formic acid (v/v), at a flow rate of 0.3 mL/min and a voltage of 4.0 kV.
